# Effects of Body Mass on Leg and Vertical Stiffness in Running Humans

**DOI:** 10.1101/2022.05.04.490639

**Authors:** Maria C. Fox, Elizabeth T. Hsiao-Wecksler, John D. Polk

**Author notes:** Corresponding author *Email address:* (John D. Polk).

## Abstract

Numerous cross-species comparisons have examined the scaling of gait parameters with respect to body mass (i.e., allometry), but few have done so within humans. This study examined how leg and vertical stiffness, force, displacement, and leg spring angle scaled in 64 healthy adults of varying body masses during slow and fast leg-length-adjusted running speeds. We calculated scaling patterns for stiffness and its components via kinematic and kinetic data using log-log regressions with 95% confidence/highest density intervals. To determine if the chosen statistical method influenced conclusions about scaling patterns, we compared regression results across three statistical methods, ordinary least squares (OLS) regression, linear mixed models (LMM), and Bayesian linear mixed models (BLMM). We also performed sex-specific analyses to determine if each sex revealed similar scaling patterns as the pooled sample. In the pooled sample, all variables scaled according to the isometric expectations, suggesting that different-sized humans move in a similar manner. Sex-specific analyses revealed similar patterns of isometry in all variables, except for vertical stiffness, which displayed slight negative allometry (i.e., lower than expected stiffness) in both sexes at the slow speed and negative allometry in females during fast running. Model choice did not significantly affect results, and scaling patterns were the same regardless of the statistical method employed.

## Introduction

### 1.1 Allometry and scaling

Body size is an important determinant of variation in limb posture and the musculoskeletal mechanisms used to resist gravity (Biewener, 1982, 1989a,b, 1990, 2005; Galilei, 1638; McMahon, 1973, 1975; Polk, 2002). Allometry describes how a given trait changes (scales) with body mass or other relevant size variable (Alexander, 1985; Gould, 1966) with allometric scaling patterns derived from linear regression equations of trait (*y*) versus size (*x*), where:

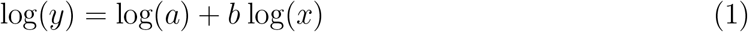

Equation (1) can also be expressed as log(*y*) ∝ *b* log(*x*) (where log(*a*), the intercept, is dropped), and finally as *y ∝ x*^*b*^, where ∝ means “proportional to” (Alexander, 1985). *b* is the allometric scaling exponent represented as the slope of the line in the log-log regression. When body mass is the size variable of interest, the scaling exponents are typically expressed as ∝ *M*^*b*^, where M is body mass.

There are different expectations for the calculated scaling exponents (*b*) based on different geometric theories. The simplest, geometric similarity, assumes that an object scales uniformly (Alexander, 1985; Hill, 1950; McMahon, 1975). According to the assumptions of geometric similarity, length (and diameter) variables should therefore scale proportional to (∝) *M* ^1*/*3^ and areas scale ∝ *M* ^2*/*3^. If traits scale uniformly according to the aforementioned expectations, this relationship is considered isometry. Deviations above this expectation (i.e., more positive/steeper slopes) are called positive allometry and deviations below this expectation (i.e., more negative/shallower slopes) are called negative allometry.

Another theory, termed dynamic similarity, was introduced by Alexander and Jayes (1983), which proposed that geometrically similar animals can also be elastically similar (i.e. deform under their body weight in a similar manner) *during movement* ; i.e., differently-sized animals would have similar kinematic parameters such as joint excursion angles (∝ *M*^0^) when moving at the same relative speed (such as a Froude speed, which is calculated as a function of leg length, gravity, and velocity: *Fr* = *V* ^2^*/gL*) (Alexander and Jayes, 1983; Alexander, 1989; Donelan and Kram, 2000). Farley et al. (1993) added that dynamically similar animals should have equal dimensionless (i.e., ∝ *M* ^0^) leg compression (Δ*L/L*_0_, where *L*_0_ is leg length) and equal dimensionless force in the leg spring (*F*_*leg*_*/BW*, where *BW* is body weight) when running at the same relative speed.

### 1.2 Leg and vertical stiffness

Limb function during running is often modeled as a spring-mass system, where the leg acts as a spring by storing and releasing elastic energy (Blickhan, 1989; Farley et al., 1991; McMahon and Cheng, 1990). In this model, leg spring stiffness is a function of force relative to the leg spring displacement, which passes through an inverted pendular arc during motion (Blickhan, 1989; Cavagna et al., 1988; McMahon and Cheng, 1990) (Figure 1). Leg spring stiffness (*k*_*leg*_) is calculated from the peak of the limb’s resultant force in the direction of the leg spring (*F*_*leg*_) divided by leg compression (Δ*L*) during the first half of stance phase (*k*_*leg*_ = *F*_*leg*_*/*Δ*L*) (Coleman et al., 2012).

**Figure 1:**
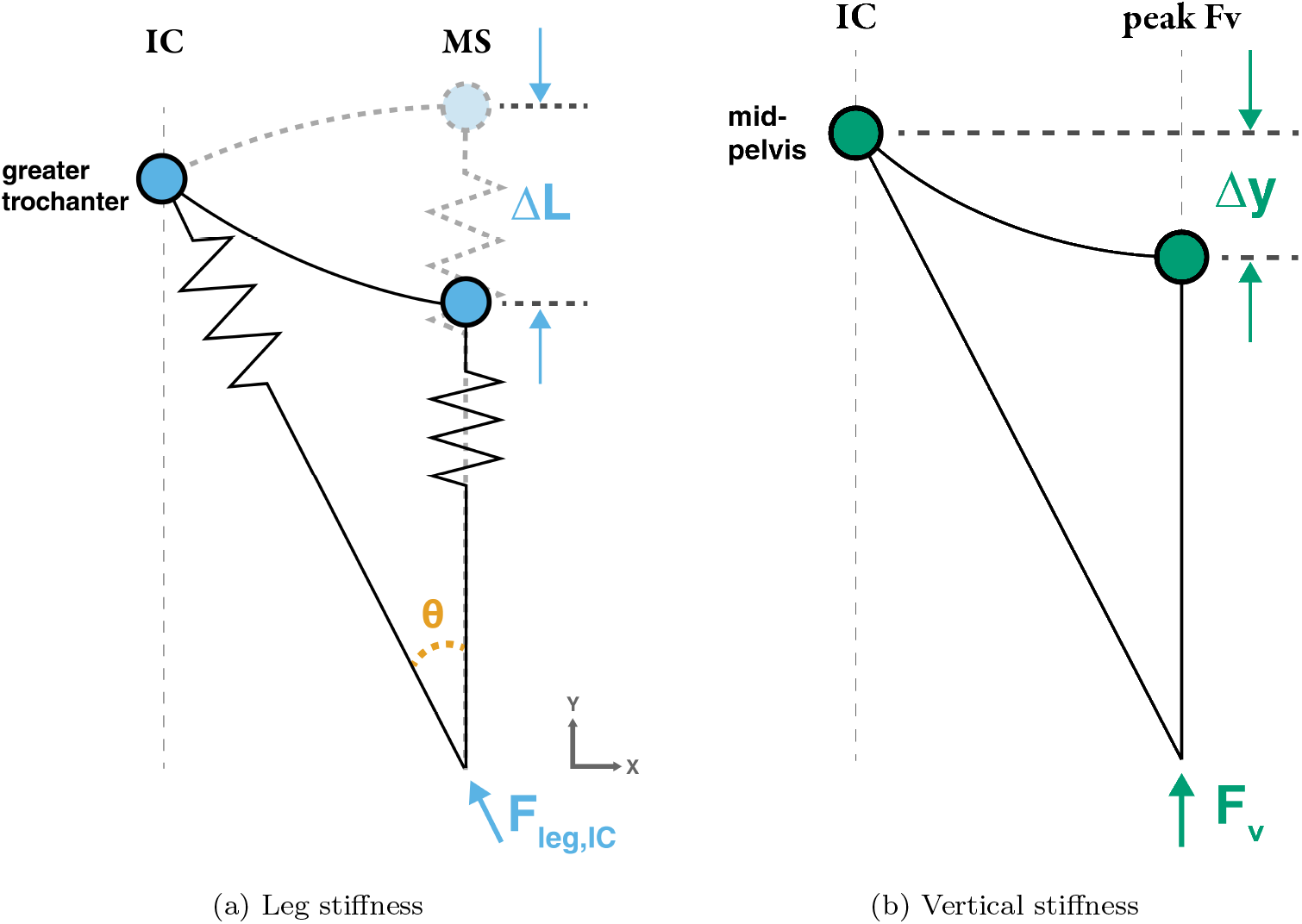
a) Leg stiffness is derived from the resultant force of the leg spring (*F*_*leg*_) divided by the change in leg compression (Δ*L*) between initial contact (IC) and mid-stance (MS). Half the angle swept by the leg spring (*θ*) is also calculated between these time points. b) Vertical stiffness is derived from the peak vertical GRF (*F*_*V*_) divided by the vertical change in the center of mass (Δ*y*) between IC and the occurrence of the peak *F*_*V*_.

Half the angle swept by the leg spring (*θ*) also represents the angular excursion of the leg spring during the first half of stance phase (Figure 1a).

Another measure of interest, vertical stiffness (*k*_*vert*_), relates peak vertical ground reaction force (*F*_*V*_) to peak vertical COM fluctuation during stance (Δ*y*) (*k*_*vert*_ = *F*_*V*_ */*Δ*y*) (Farley 2 et al., 1993). Vertical stiffness can be used to infer the frequency of vertical motions during running and is directly related to contact time during running (Brughelli and Cronin, 2008; Cavagna et al., 1988; Farley et al., 1993; McMahon et al., 1987).

### 1.3 Scaling of stiffness

Lower limb stiffness (*k*_*leg*_, *k*_*vert*_) and its components of force (*F*_*leg*_, *F*_*V*_) and displacement (Δ*L*,Δ*y*) are an interesting natural test case for dynamic similarity, since this theory applies to bodies in movement. Scaling expectations adhering to dynamic similarity (i.e., geometric similarity at equal relative speeds) are reported in Table 1. Leg spring angle (*θ*) is predicted by dynamic similarity to be invariant across body size and therefore does not technically “scale”.

**Table 1:** Scaling expectations according to dynamic similarity

| Variable | Isometry |
| --- | --- |
| Force ($kN$) | $M^1$ |
| Displacement ($m$) | $M^{1/3}$ |
| Stiffness ($kN/m$) | $M^{2/3}$ |
| Leg angle ($\theta$, $^\circ$) | $M^0$ |

Previous work found scaling exponents of *M* ^0.67^ for leg stiffness and *M* ^0.61^ for vertical stiffness in interspecific samples of quadrupedal mammals (mass between 0.1 and 135 kg) (Farley et al., 1993). Herr (1998), using simulated model data, reported that leg stiffness scaled ∝*M* ^0.69^ and vertical stiffness ∝*M* ^0.61^ in quadrupeds, which were almost identical to the values reported by Farley et al. (1993). Larger mammals therefore appear to have stiffer leg springs that undergo less vertical displacement, as predicted by geometric similarity. Farley et al. (1993) observed that peak *F*_*leg*_ scaled ∝*M* ^0.97^ and Δ*L* scaled ∝*M* ^0.30^, all close to the geometric similarity expectation. Farley et al. (1993) also reported that leg spring angle (*θ*) scaled ∝*M* ^−0.03^, Δ*L/L*_0_ scaled ∝*M* ^−0.04^, and *F*_*leg*_*/BW* scaled ∝*M* ^−0.03^, which suggests that the animals in their study conformed to dynamic similarity.

Only two studies have examined how stiffness scales with body size in humans (Carruthers and Farley, 1998; Farley and Korff, 1999). Carruthers and Farley (1998) reported that peak force scaled ∝*M* ^0.87^, leg compression ∝*M* ^0.21^, and leg stiffness ∝*M* ^0.66^ among a sample of 21 adults (sex not reported) running at 4.0 m/s. This study concluded that stiffness scaled “steeply” (Carruthers and Farley, 1998, p307) with body mass, but these results were very close to the isometric expectations. Farley and Korff (1999) also examined leg stiffness during hopping in 18 adults; they reported much higher scaling exponents (∝*M* ^1.00*±*0.23^). This higher value is likely due to considerably smaller leg compression during the hopping task; reported values for leg compression during running were approximately twice as high as those during hopping in these studies (Carruthers and Farley, 1998; Farley et al., 1993; Farley and Korff, 1999).

There are multiple reasons to question the applicability of geometric similarity in humans given the wide range of variation in human body and limb proportions (Nevill et al., 2004; Steudel-Numbers and Weaver, 2006; Sylvester et al., 2008; Kramer and Sylvester, 2013). Although humans may not be perfectly isometric in their limb lengths, it is unclear whether these proportional differences are enough to violate geometric similarity (Fox et al., 2021).

### 1.4 Current study

The current study had two main goals. The first was to evaluate how leg and vertical stiffness (and their determinants) scale in a large sample of humans, which would enable empirical evaluation of whether intraspecific (within-species) scaling of stiffness differs from interspecific expectations and whether humans conform to dynamic similarity. We hypothesized that humans would conform to dynamic similarity and display similar scaling patterns as other animals (Table 1). The second goal was to evaluate sex-specific scaling of stiffness properties, which would determine if male and female runners differ in running behavior and whether sex-specific scaling patterns match those seen in sex-pooled samples.

## 2. Methods

### 2.1 Subjects

64 healthy (*BMI <* 27) females (N=35) and males (N=29) aged 18-35 with no history of gait-related pathologies or recent lower limb injuries (*<*1yr) were recruited to participate in this study. Most subjects were recreationally active, but experienced runners were not specifically recruited for this study. All protocols were approved by the University of Illinois IRB and subjects gave written informed consent for participation.

Subjects were recruited primarily to maximize variation in height across both sexes. We did so by recruiting relatively short and tall individuals for each sex who were below ≈25% (F ≤ 158 cm, M ≤ 170 cm) and above ≈75% (F ≥ 167 cm, M ≥ 178 cm) of the reported average adult heights in the United States (Fryar et al., 2012). Due to using a population of convenience of predominantly recreationally active young adults, subjects also had a relatively even distribution of body mass and BMI across the desired height ranges.

### 2.2 Measurements

We collected anthropometric data for each subject to describe overall body size and shape. Mass was calculated using a force plate (AMTI, Watertown, MA). Running trials were performed on a split-belt Bertec instrumented treadmill (Bertec, Columbus, OH). Subjects wore their own athletic/running shoes for data collection. All subjects wore spandex shorts, and females wore a sports bra. Motion capture data were collected with a 6-camera Qualisys system (Qualisys AB, Gö teborg, Sweden) and 48 reflective markers adhered to skin or clothing/shoes. Markers were positioned on common anatomical landmarks and segmental triads (Table A1).

Before running trials began, subjects acclimated to the treadmill by walking at several different speeds. Because not all subjects were experienced runners, subjects were secured via a custom nylon harness attached to a moving trolley during running trials in the event of a fall. Subjects practiced running with the harness before trials were recorded to acclimate the subject to the harness. A single-axis load cell (LCM200, Futek, Irvine, CA) was attached to the top of the harness to verify that the harness was not being loaded during running trials. Subjects ran on one belt of the treadmill at a time and switched belts each trial to avoid overheating either belt.

Data were collected at two different running speeds determined by Froude number (*Fr* = *V* ^2^*/gL, V* =speed in m/s, *g* =9.81 *m/s*^2^, *L* = leg length in m):

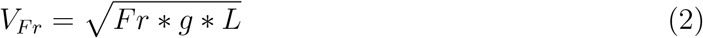

The slower speed at the smaller Froude number (*Fr* = 0.8, N=64) was approximately twice as fast as a comfortable walking speed and equated to a slow jog (mean 2.6 *±* 0.1 m/s). The faster speed at the higher Froude number (*Fr* = 1.8) was approximately three times as fast as walking and resembled an endurance running pace (mean 4.0 *±* 0.1 m/s) (Bramble and Lieberman, 2004; Cavanagh and Kram, 1989). Two subjects opted not to perform the faster speed (N=62 for the fast speed). We refer to the aforementioned speeds as slow and fast Froude speeds for the remainder of this article. Three trials were collected at each running speed (6 trials total, 3 per speed) with a short break in between each trial. Slow Froude trials lasted 15s each and fast Froude trials lasted 10s each. Trial recordings began after the treadmill reached the desired speed.

Leg and vertical stiffness (*k*_*leg*_ and *k*_*vert*_, respectively) and their force (*F*_*leg*_ and *F*_*V*_) and displacement (Δ*L* and Δ*y*) components were computed for each gait cycle (defined by the period during which the vertical force threshold exceeded 20N) using direct kinematic and kinetic data (Coleman et al., 2012) (see detailed calculations in (Fox, 2020)). Unrealistic values (e.g., stiffness *<* 0) were removed from the analysis.

‘Direct’ methods to calculate stiffness utilize 3D motion capture and force data to calculate each parameter, and are therefore arguably more precise than estimation-based methods Coleman et al. (2012). In this study, COM position was approximated by averaging the four pelvis marker locations, as opposed to the alternate approach of double integrating the vertical acceleration of force (Cavagna, 1975). Half the angle swept by the leg spring (*θ*) was also calculated using kinematic data. All data were processed in MATLAB R2018a (MathWorks, Natick, MA).

Dimensionless metrics can account for size effects (Farley et al., 1993; Hof, 1996). In theory, dimensionless variables should be invariant across body size (∝*M* ^0^) if the animals are dynamically similar. Dimensionless variables were also included in analyses, and were calculated for all variables (except for *θ*) using the equations found in (Farley and González, 1996; McMahon and Cheng, 1990).

### 2.3 Statistical analysis

#### 2.3.1. Scaling of leg and vertical stiffness

Leg stiffness (*k*_*leg*_) and its components (*F*_*leg*_ and Δ*L*), vertical stiffness (*k*_*vert*_) and its components (*F*_*V*_ and Δ*y*), leg spring angle (*θ*), and dimensionless variables were the variables of interest presented in this study. All statistical analyses were performed in R v3.6.0-3.6.2 (R Core Team, 2019).

Allometric scaling exponents were calculated for each variable using the log-log regression expression presented in Equation 1 as a function of body mass. Model fit statistics and effect size measures are reported for each model and analysis. Separate models were used for the two Froude speeds to account for variation in Froude number.

We tested each variable with three different statistical models to determine whether the slopes (scaling exponents) and their associated confidence intervals were influenced by model choice. The first model, ordinary least squares linear regression (OLS), used aggregated data for each subject and condition.

The second model incorporated a random effect (intercept) for subjects using a linear mixed model (LMM) with the non-aggregated repeated measures data. LMMs incorporate both fixed effects (e.g., a condition such as running speed or sex) and random effects, which are uncontrolled, unobserved “random” variables (e.g., subject). LMM models allow users to partition the error variance in the model according to those random effects as well as the residual error of the model.

The final model represented a Bayesian extension of the linear mixed model (BLMM). BLMMs apply Bayes’ theorem to regression modeling, where prior beliefs about the parameter(s) of interest are used to inform posterior beliefs about the parameter(s) of interest given the observed data (Kruschke, 2014). BLMMs report model coefficients as posterior probability distributions, which represent “a compromise between the data model (likelihood) and the prior, and describes the relative plausibility of all parameter values conditional on the model” (Muth et al., 2018, p. 101). BLMMs were performed using the rstanarm R package (Goodrich et al., 2020). We fitted BLMMs (estimated using Markov Chain Monte Carlo (MCMC) sampling with 6 chains of 20,000 iterations, 1000 warmups, and a thinning interval of 10) with subject as the random effect.

For each model, equivalent model fit was calculated using *R*^2^ and uncertainty interval (Table 2). For OLS and LMM models, uncertainty level was assessed by a 95% confidence interval. For the BLMM model, uncertainty level was assessed by a 95% posterior highest density interval (HDI). If the 95% uncertainty interval of the scaling exponent (*b*) fell outside the isometric expectation, then we concluded that positive or negative allometry was present.

**Table 2:** Statistical models and parameters

| Model | R Package | Slope | Uncertainty Interval | $R^2$ |
| --- | --- | --- | --- | --- |
| OLS<sup>1</sup> | `lmodel2` | Estimate | 95% CI | Traditional $R^2$ |
| LMM<sup>2</sup> | `lme4` | Fixed Effect Estimate | 95% CI | Conditional $R^2$ <sup>3</sup> |
| BLMM<sup>4</sup> | `rstanarm` | Posterior Median | 95% HDI | Bayesian $R^2$ <sup>5</sup> |

<sup>1</sup> Legendre (2018)
<sup>2</sup> Bates et al. (2014)
<sup>3</sup> $R^2$ value taking both the fixed and random effects into account (Nakagawa and Schielzeth, 2013; Nakagawa et al., 2017)
<sup>4</sup> Goodrich et al. (2020)
<sup>5</sup> Median of Bayesian approximation of conditional $R^2$ (Gelman et al., 2019)

#### 2.3.2. Sex differences

To determine whether sex differences were present in the slopes of each variable, we ran Bayesian linear mixed models incorporating an interaction effect between mass and sex. We chose to use BLMMs for this analysis because they provided the most stable model coefficients and had similar results to the other statistical methods. We then computed estimated marginal trends using the emmeans R package (Lenth, 2019), which produced sexspecific slopes. We examined upper and lower 95% HDIs from each sex-specific slope to determine if positive or negative allometry was present within each sex. To determine if sex-specific slopes differed from one another (Female-Male contrast), we examined pairwise comparisons of the estimated marginal trends, which provided a median estimate of the difference between the slopes as well as a 95% HDI of that difference. If the 95% difference HDI (95% Diff HDI) contained zero, the slopes were not significantly different from one another.

## 3. Results

### 3.1. Scaling of leg and vertical stiffness

Subjects in this study spanned ≈ 50*cm* in height, ≈ 35*cm* in leg length, and ≈ 50*kg* in mass (Table 3). Due to variations in leg length, speeds at the slow Froude No. ranged from ≈2.4-2.9m/s; speeds at the fast Froude No. ranged from ≈ 3.4-4.4m/s.

**Table 3:** Descriptive statistics for subjects split by sex.

|  | Overall<br>N=64 |  |  | Female<br>N=35 |  |  | Male<br>N=29 |  |  |
| --- | --- | --- | --- | --- | --- | --- | --- | --- | --- |
| | Mean ± SD | Min | Max | Mean ± SD | Min | Max | Mean ± SD | Min | Max |
| Age (*yrs*) | 22.2 ± 3.8 | 18.0 | 33.0 | 22 ± 3.4 | 18.0 | 30.0 | 22.3 ± 4.4 | 18.0 | 33.0 |
| BMI (*kg/m<sup>2</sup>*) | 22.6 ± 2.3 | 18.0 | 26.7 | 22.6 ± 2.3 | 18.0 | 26.5 | 22.5 ± 2.3 | 18.0 | 26.7 |
| Height (*cm*) | 168.6 ± 13.4 | 141.5 | 191.8 | 162.2 ± 11.5 | 141.5 | 181.0 | 176.4 ± 11.3 | 153.5 | 191.8 |
| Leg Length (*cm*) | 88.0 ± 8.6 | 70.1 | 106.0 | 84.2 ± 7.7 | 70.1 | 98.4 | 92.5 ± 7.4 | 79.9 | 106.0 |
| Mass (*kg*) | 63.8 ± 11.2 | 41.1 | 89.9 | 59.1 ± 9.4 | 41.1 | 78.2 | 69.6 ± 10.7 | 47.2 | 89.9 |

The OLS, LMM, and BLMM models did not differ appreciably in reported scaling exponents or 95% intervals (CI or HDI; Figures 2a, 3a, and 4A; Table A2). While the scaling exponents and 95% intervals did not differ across models, explained variance (*R*^2^) was considerably higher for models containing random effects (LMM and BLMM). For example, *R*^2^ values for Δ*L* during slow running trials were 0.32 (OLS), 0.87 (LMM), and 0.86 (BLMM) (Table A2).

**Figure 2:**
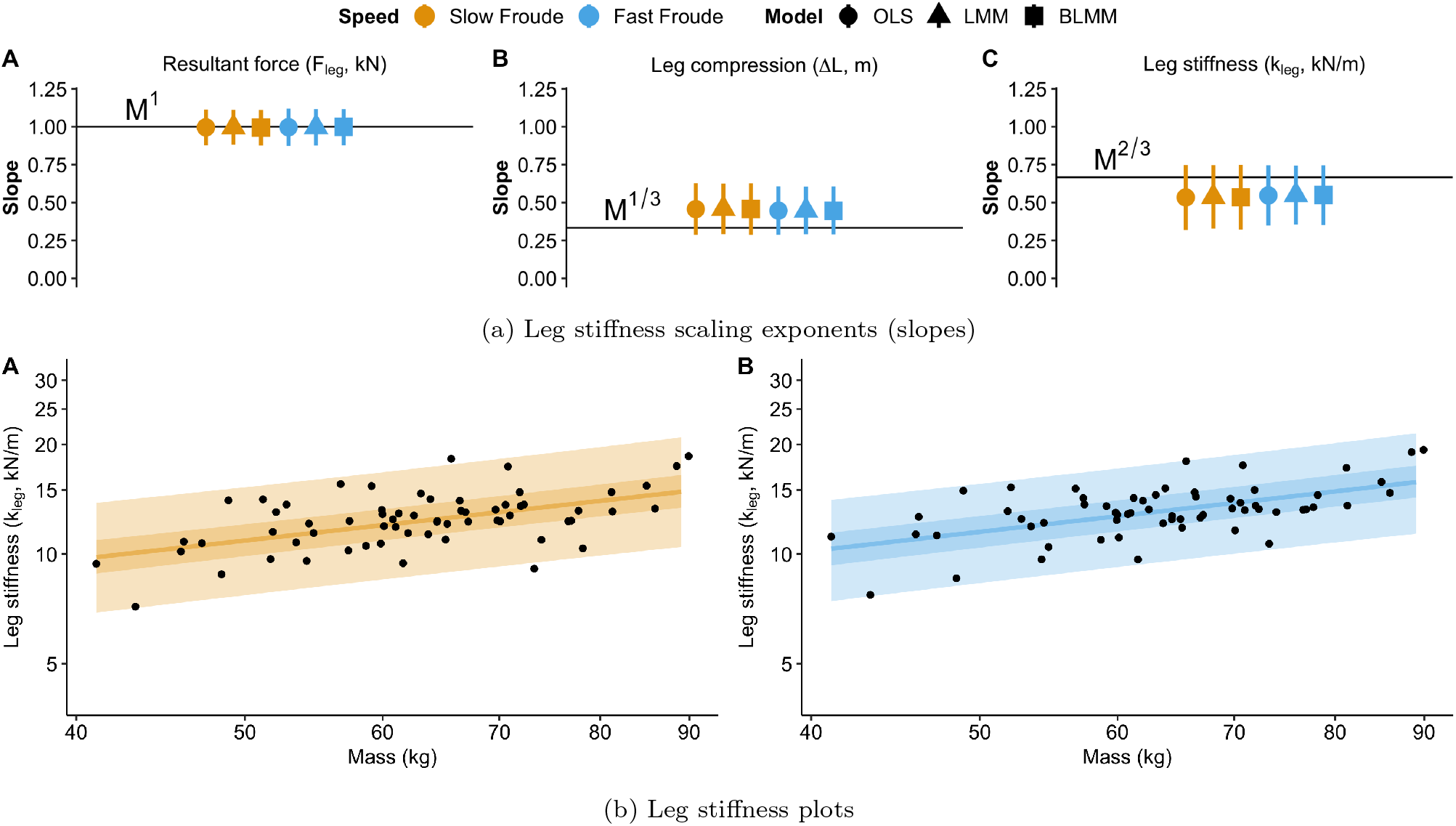
(a) Top figure displays the slope estimate and 95% CI/HDI for each model. (b) Bottom figure displays leg stiffness values at (A) slow and (B) fast running speeds plotted on a log-log scale with 50% (darker) and 95% (lighter) Bayesian posterior prediction intervals from BLMM model.

**Figure 3:**
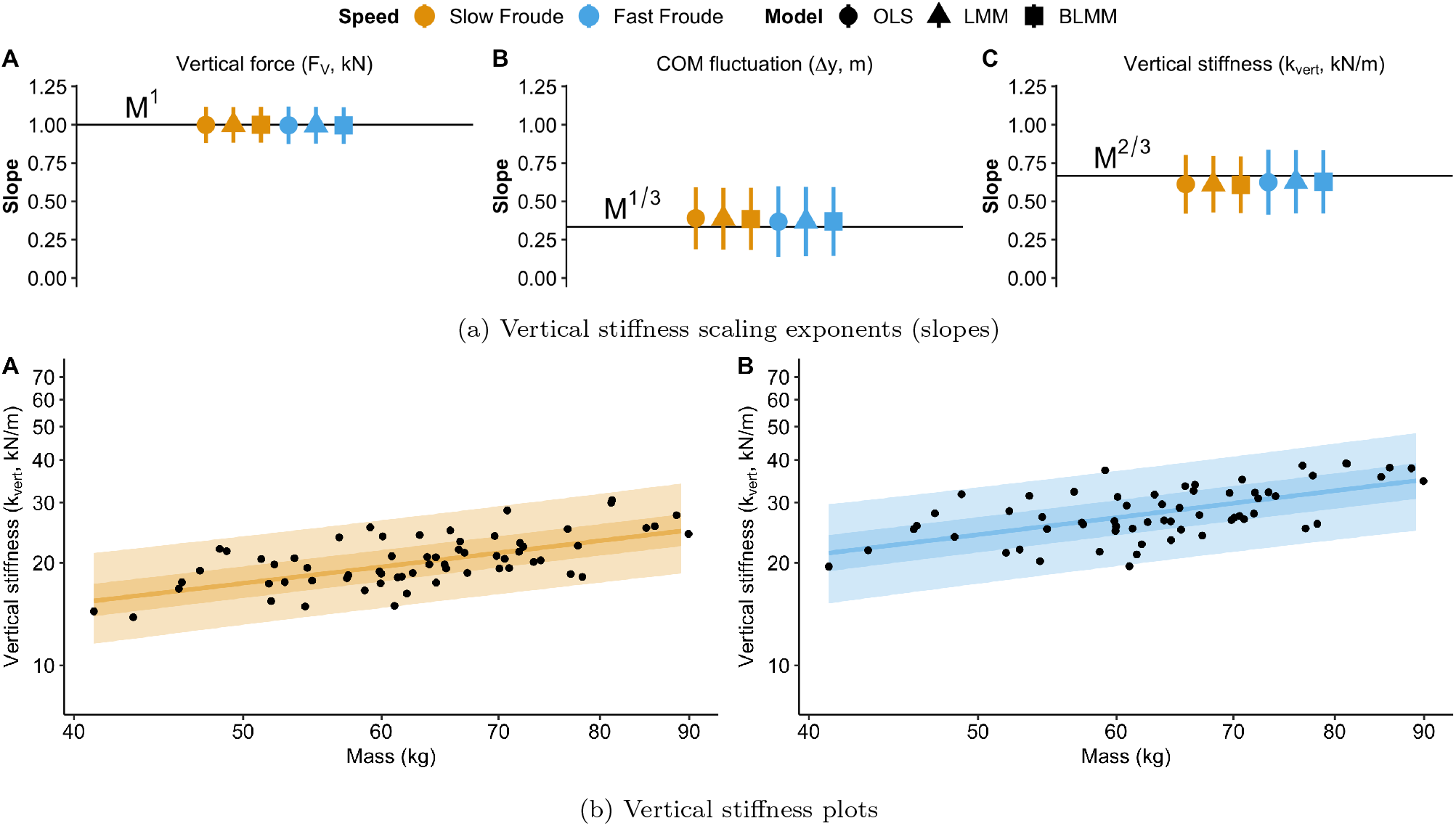
(a) Top figure displays the slope estimate and 95% CI/HDI for each model. (b) Bottom figure displays vertical stiffness values at (A) slow and (B) fast running speeds plotted on a log-log scale with 50% (darker) and 95% (lighter) Bayesian posterior prediction intervals from BLMM model.

Average scaling exponents for *θ* were invariant across body size and scaled ∝ *M* ^0.04^ during slow running and ∝ *M* ^0.05^ during fast running (Figure 4).

**Figure 4:**
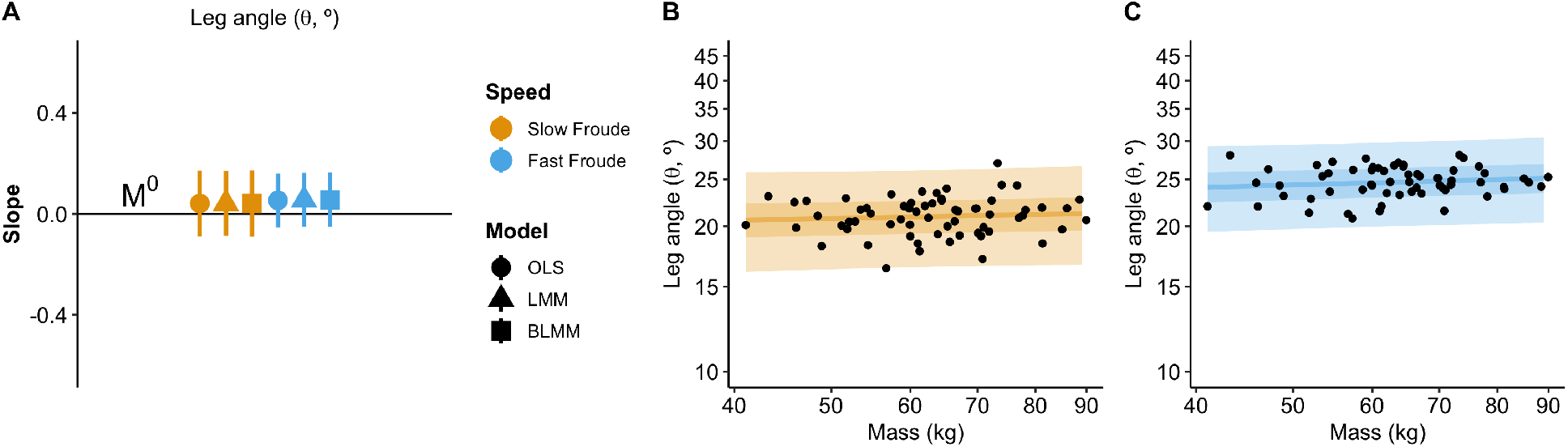
(A) Slope estimates and 95% CI/HDI for each leg spring angle (*θ*) model. *θ* values at (B) slow and (C) fast running speeds plotted on a log-log scale with 50% (darker) and 95% (lighter) Bayesian posterior prediction intervals from BLMM model.

Averaged across the three statistical models, leg stiffness and its components increased with body mass: *k*_*leg*_ scaled ∝ *M* ^0.53^ (slow) and ∝*M* ^0.55^ (fast), *F*_*leg*_ scaled ∝ *M* ^1.00^ (slow) and ∝ *M* ^1.00^ (fast); and Δ*L* scaled ∝ *M* ^0.46^ (slow) and ∝ *M* ^0.45^ (fast) (Figure 2a). Leg compression exponents approached positive allometry (*> M* ^1*/*3^) and leg stiffness trended toward negative allometry (*< M* ^2*/*3^) at both speeds in this sample.

Averaged across the three models, vertical stiffness and its components also increased with body mass: *k*_*vert*_ scaled ∝ *M* ^0.61^ (slow) and ∝ *M* ^0.63^ (fast); *F*_*V*_ scaled ∝ *M* ^1.00^ (slow) and ∝ *M* ^1.00^ (fast); and Δ*y* scaled ∝ *M* ^0.39^ (slow) and *M* ^0.37^ (fast) (Figure 3a). Vertical stiffness and its component scaling exponents adhered to isometric expectations at both speeds in this sample. Vertical stiffness displayed higher variability at the fast speed (as observed by the width of the 95% posterior prediction interval in Figure 3b).

Dimensionless (size-corrected) variables revealed scaling exponents that were invariant across body mass (Figure 5, Table A4). Averaged across models, dimensionless leg stiffness (*K*_*LEG*_) scaled ∝ *M* ^−0.03^ (slow) and ∝ *M* ^−0.01^ (fast). Dimensionless vertical stiffness (*K*_*V ERT*_) scaled ∝ *M* ^0.05^ (slow) and ∝ *M* ^0.07^ (fast). Dimensionless force variables (*F*_*leg*_ and *F*_*V dim*_) as well as dimensionless leg compression (Δ*L*_*dim*_) and COM fluctuation (Δ*y*_*dim*_) were invariant across body size at both speeds (Table A4). Confidence intervals/HDIs for every dimensionless variable and speed included zero (Table A4, Table A5).

**Figure 5:**
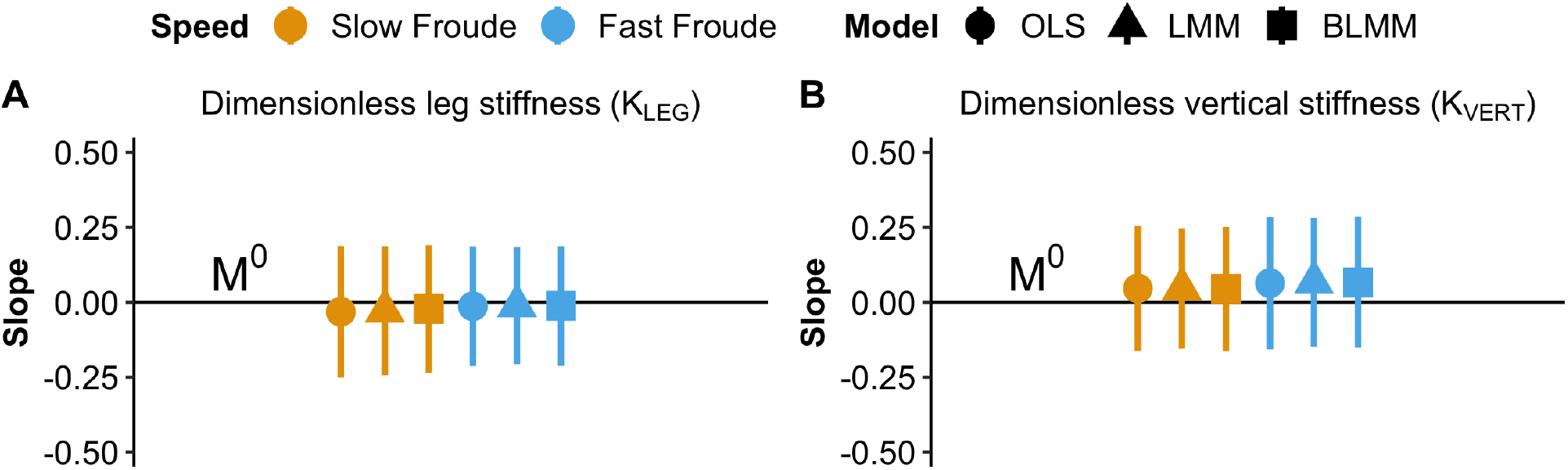
Dimensionless slope estimates and 95% CI/HDI for each model and speed for (A) *K*_*LEG*_ and (B) *K*_*V ERT*_.

### 3.3. Sex differences

Vertical stiffness was the only variable that contained HDIs outside the isometric expectations in the sex-specific analysis (Table 4). Females and males displayed slight negative allometry (∝ *M* ^0.39^, ∝ *M* ^0.45^, respectively) during slow running; females displayed slight negative allometry during fast running (∝ *M* ^0.41^). These effects were not observed when sexes were pooled.

**Table 4:** Sex comparison model results. Diff Est is the median slope difference (contrast) between females and males, and Diff HDI is the 95% HDI interval of that difference. * denotes that slope confidence interval lies outside the isometric expectation.

| Variable | Speed | Sex | Slope Est | 95% HDI | $R^2$ | Diff Est | 95% Diff HDI |
| --- | --- | --- | --- | --- | --- | --- | --- |
| $F_{leg}$ | Slow Froude | Female | 0.94 | 0.79, 1.09 | 0.97 | 0.02 | -0.14, 0.18 |
|  |  | Male | 0.92 | 0.77, 1.08 |  |  |  |
|  | Fast Froude | Female | 0.90 | 0.75, 1.04 | 0.97 | -0.01 | -0.17, 0.14 |
|  |  | Male | 0.91 | 0.76, 1.06 |  |  |  |
| $\Delta L$ | Slow Froude | Female | 0.50 | 0.30, 0.71 | 0.86 | 0.05 | -0.10, 0.20 |
|  |  | Male | 0.45 | 0.25, 0.66 |  |  |  |
|  | Fast Froude | Female | 0.41 | 0.22, 0.59 | 0.84 | 0.05 | -0.09, 0.20 |
|  |  | Male | 0.36 | 0.18, 0.55 |  |  |  |
| $k_{leg}$ | Slow Froude | Female | 0.43 | 0.18, 0.68 | 0.88 | -0.05 | -0.23, 0.14 |
|  |  | Male | 0.47 | 0.20, 0.72 |  |  |  |
|  | Fast Froude | Female | 0.49 | 0.26, 0.72 | 0.85 | -0.06 | -0.25, 0.10 |
|  |  | Male | 0.55 | 0.32, 0.79 |  |  |  |
| $F_V$ | Slow Froude | Female | 0.94 | 0.79, 1.09 | 0.97 | 0.02 | -0.14, 0.18 |
|  |  | Male | 0.92 | 0.77, 1.08 |  |  |  |
|  | Fast Froude | Female | 0.90 | 0.75, 1.04 | 0.97 | -0.01 | -0.17, 0.15 |
|  |  | Male | 0.91 | 0.76, 1.06 |  |  |  |
| $\Delta y$ | Slow Froude | Female | 0.54 | 0.30, 0.77 | 0.77 | 0.05 | -0.12, 0.22 |
|  |  | Male | 0.49 | 0.25, 0.73 |  |  |  |
|  | Fast Froude | Female | 0.47 | 0.21, 0.76 | 0.80 | 0.06 | -0.12, 0.25 |
|  |  | Male | 0.41 | 0.14, 0.68 |  |  |  |
| $k_{vert}$ | Slow Froude | Female | 0.39* | 0.20, 0.60 | 0.82 | -0.06 | -0.22, 0.11 |
|  |  | Male | 0.45* | 0.24, 0.65 |  |  |  |
|  | Fast Froude | Female | 0.41* | 0.18, 0.64 | 0.82 | -0.10 | -0.28, 0.09 |
|  |  | Male | 0.51 | 0.28, 0.75 |  |  |  |
| $\theta$ | Slow Froude | Female | 0.05 | -0.11, 0.21 | 0.61 | 0.00 | -0.10, 0.12 |
|  |  | Male | 0.05 | -0.11, 0.20 |  |  |  |
|  | Fast Froude | Female | 0.04 | -0.08, 0.18 | 0.54 | 0.01 | -0.09, 0.11 |
|  |  | Male | 0.03 | -0.10, 0.17 |  |  |  |

The sex-specific slopes (scaling exponents) were smaller (shallower) for *F*_*leg*_, *F*_*V*_, *k*_*leg*_, and *k*_*vert*_ at both speeds, whereas they were larger (steeper) for Δ*L* and Δ*y* when compared to the pooled results (Table 4 and A2). This difference was likely due to influential observations at the tails (i.e., low and high body mass) that affect the regressions differently at the pooled and sex-specific levels.

Contrasts of estimated marginal trends revealed no statistically credible differences in the slopes of males and females (Table 4). These results were confirmed by the wide HDIs of the differences, all of which encompassed zero (Table 4, Table A6).

## 4. Discussion

Due to the relatively wide variation in the data, none of the statistical models examined in this study excluded the isometric expectations (Table 1) for pooled sexes, therefore model choice had no effect on the results. This result suggests that humans appear to conform to expectations of dynamic similarity in measures of force, displacement, stiffness, and leg spring angle when sexes are pooled. Dimensionless variables were also invariant across body mass (∝ *M* ^0^).

Leg compression trended toward positive allometry in the pooled sample, accompanied by a trend towards negative allometry in leg stiffness. These results suggest that larger individuals may have relatively greater leg compression and subsequently less stiff leg springs than expected for their mass.

Confidence intervals/HDIs for all dimensionless variables encompassed the *M* ^0^ expectation. These results suggest that humans experience equal ground reaction forces relative to their body weight and equal levels of leg compression relative to their leg lengths. When leg length and mass are both accounted for, stiffness (leg and vertical) is invariant across body sizes.

Dimensionless leg stiffness values in our sample closely match those reported by Blickhan and Full (1993) and Shen and Seipel (2015a,b, 2018), who state that dimensionless stiffness values in this range represent both an energetic optimum and maximum locomotor stability (slow Froude averaged subject range *K*_*LEG*_ = 11.77 − 24.25; fast Froude averaged subject range *K*_*LEG*_ = 12.04 − 23.90). The results of this study suggest that despite variability in multiple factors (e.g., body proportions, running behavior, and/or training), intraspecific patterns of stiffness in humans are both dynamically similar and may represent an energetic optimum. Rathkey and Wall-Scheffler (2017) observed that males choose to run at speeds close to their energetic optimum (i.e., speed at which cost of transport is minimized). Their preferred speeds corresponded to a mean Froude number of 1.74, which was similar to our fast Froude speed (1.8).

### 4.1. Sex differences

There were no significant interactions between sexes in the models presented here, and slope estimates were similar between the sexes. When split by sex, both sexes were outside the isometric expectation (∝ *M* ^2*/*3^) for vertical stiffness during slow running, displaying slight negative allometry (Table 4). Additionally, females displayed negative allometry during fast running. In contrast, vertical stiffness with pooled sexes did not show this effect (Figure 3a).

### 4.2. Study limitations

There are multiple limitations to this study. In a study on geometric similarity of anthropometric dimensions that included the sample presented here, we determined that sexspecific regressions only departed from isometry in one dimension tested (shoulder width) (FOX submitted). However, BMI was constrained in the current sample (18-27 *kg/m*^2^). It is possible that a wider range of BMIs would have accompanied larger departures from geometric similarity and therefore violate dynamic similarity. A future study including an even wider range of BMI (i.e., BMI *>* 30), mass, and height might present different results.

There are a number of other external factors that might affect leg and vertical stiffness that were not examined or controlled for in this study. While surface stiffness was held constant in this study, it is unclear if we would observe the same patterns on another surface, since research indicates that runners adjust their limb stiffness on different surfaces (Ferris et al., 1998, 1999; Kerdok et al., 2002).

Research on shoe cushioning (i.e., minimal shoes, conventional shoes, and/or maximally-cushioned shoes) indicates that runners modulate their leg stiffness to manage impact loading and potentially reduce injuries (Bishop et al., 2006; Divert et al., 2005; Kulmala et al., 2018; Lussiana et al., 2015). Subjects wore their own shoes during the study, so it is also possible that shoe cushioning affected variability and/or values of stiffness. Anatomical variability in foot arch height has also been shown to affect values of leg stiffness (Williams et al., 2004). It is also possible that foot strike patterns (rearfoot vs. forefoot/midfoot strikes) influenced stiffness (Butler et al., 2003; Hamill et al., 2014; Seyfarth et al., 2002), and a future analysis of these data could quantify foot strike patterns and compare stiffness among subjects using different strike patterns.

## 5. Conclusions

We observed that across sexes, humans conform to isometric expectations for dynamic similarity for all variables considered, suggesting that humans follow similar patterns as other animals for stiffness (∝ *M*^2*/*3^), force (∝ *M*^1^), displacement (∝ *M*^1*/*3^), and leg spring angle (∝ *M* ^0^). There was a slight (but insignificant) trend towards negative allometry in leg stiffness (*< M* ^2*/*3^), suggesting that, perhaps with a larger and more extreme BMI range or larger sample size, heavier humans may have relatively higher leg compression than expected for mass. Statistical model choice had little to no influence on the results of the scaling analysis.

These same patterns held true with sex-specific analyses, except for slight negative allometry observed in vertical stiffness in both males and females during slow running, and in females during fast running. All metrics displayed considerable variability among subjects, but despite this variability, the individuals in this sample conformed to dynamic similarity when running at the same relative speed, as was hypothesized.

## 7. Acknowledgments

This study was supported by the Wenner-Gren Foundation, the University of Illinois Beckman Institute for Advanced Science and Technology, and the University of Illinois Department of Anthropology.

We would also like to thank all the anonymous individuals who participated in this study, research assistants Milena Singletary and Elizabeth Phillips for their help conducting the study and processing data, members of the Human Dynamics and Controls Lab for their help and support (especially Chenzhang Xiao), and Alena Grabowski for her input on the treadmill harness system.

