## Supplemental Data for "Effects of Body Mass on Leg and Vertical Stiffness in Running Humans"

### 8. Appendix

### 9. Motion capture marker setup

Table A1: Description and names of markers used with motion capture data.

| Segment | Marker Name(s) | Landmark |
|---|---|---|
| Torso | RACR, LACR<br>CLAV<br>C7 | Acromion Process<br>Mid-point of clavicles<br>C7 spinous process |
| Upper Arm | RBICEPU, LBICEPU<br>RBICEP, LBICEP<br>RBICEPL, LBICEPL<br>RELBOW, LELBOW | Upper bicep<br>Midpoint of bicep<br>Lower bicep<br>Lateral epicondyle of humerus |
| Lower Arm | RFARM, LFARM<br>RRAD, LRAD<br>RULNA, LULNA | Midpoint of forearm<br>Radial styloid process<br>Ulnar styloid process |
| Pelvis | RASIS, LASIS<br>RPSIS, LPSIS | Anterior superior iliac spine<br>Posterior superior iliac spine |
| Thigh | RTHIGHU, LTHIGHU<br>RTHIGH, LTHIGH<br>RTHIGHL, LTHIGHL<br>RKNEE, LKNEE | Upper thigh<br>Midpoint of thigh<br>Lower thigh<br>Lateral epicondyle of femur |
| Shank | RSHANKU, LSHANKU<br>RSHANK, LSHANK<br>RSHANKL, LSHANKL<br>RANK, LANK | Upper shank<br>Midpoint of shank<br>Lower shank<br>Lateral malleolus of fibula |
| Foot | RHEEL, LHEEL<br>RMTI, LMTI<br>RMTV, LMTV | Calcaneal tuberosity<br>Head of first metatarsal<br>Head of fifth metatarsal |

### 10. Stiffness calculations

This method followed Coleman et al. (2012), using a direct kinematic and kinetic approach to calculate leg stiffness. We also devised an equivalent direct method to calculate vertical stiffness.

**Leg Stiffness**

$$\theta_L = tan^{-1}(\frac{A}{B}) \tag{3}$$

Equation 3 represents the angle between $L$ and the horizontal axis, where $A$ was the vertical distance between the greater trochanter and the floor, $B$ was the horizontal (AP) distance between the greater trochanter and the mid-toe position (Figure A1).[1] $L$ was the true leg length, calculated by $\sqrt{A^2 + B^2}$.

$$\theta_R = cos^{-1}(\frac{F_V}{F_R}) \tag{4}$$

Equation 4 represents the angle at which resultant force ($F_R$) was directed relative to the vertical axis, where $F_R = \sqrt{F_V^2 + F_H^2}$.

$$\theta_{leg} = (90 - \theta_L) - \theta_R \tag{5}$$

Equation 5 represents the angle (in degrees) between the line of $F_R$ and the line along which $F_{leg}$ was directed.

$F_{leg}$ = maximum resultant force calculated in the direction of the leg spring during the braking phase (initial contact to mid-stance, defined here by the point where the horizontal (AP) force component crossed zero during stance).

$$F_{leg} = max(F_R cos(\theta_{leg})) \tag{6}$$

Finally, leg stiffness is calculated as:

$$k_{leg} = \frac{F_{leg}}{\Delta L} \tag{7}$$

[1] Mid-toe position was calculated as the horizontal mid-point between the first and fifth metatarsal foot markers; note that in Coleman et al. (2012) COP is used instead. We chose to use the mid-toe position because there was considerable signal noise in the COP position that proved problematic for results of $\Delta L$. Mid-toe position tracked closely with that of the COP and therefore provided a more stable alternative than COP position.

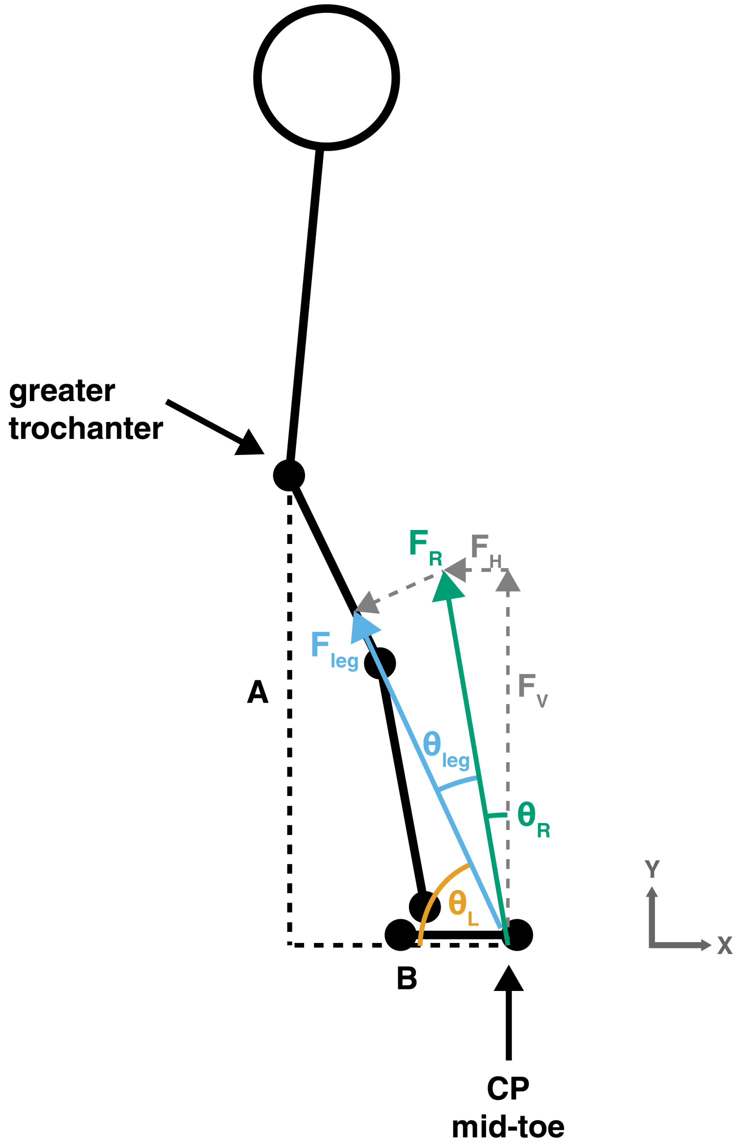


Figure A1: Illustration of direct method for calculating leg stiffness. Center of pressure (CP) was used in force calculations ($F_V$, $F_H$, and $F_R$); mid-toe position was used to calculate $B$ and $\theta_L$. Adapted from Coleman et al. (2012).

#### Vertical Stiffness

$$k_{vert} = \frac{F_V}{\Delta y} \tag{8}$$

$$\Delta y = y_{IC} - y_{min} \tag{9}$$

$F_V$ is the maximum vertical ground reaction force during stance phase (which often coincided with mid-stance). Equation 9 calculates maximum vertical fluctuation in COM during stance phase (from initial contact (IC) to the minimum vertical height of the COM, which coincided with mid-stance and the peak vGRF. COM position was approximated using the vertical mid-pelvis position, calculated as the mean point of four pelvis markers (RASIS, LASIS, RPSIS, LPSIS).

**Theta**

$\theta$ = half the angle swept by the leg spring, calculated as the difference in leg spring angle (Equation 11, in degrees) between initial contact and mid-stance. The angle is calculated using the vertical (Y) and horizontal (X) positions of the greater trochanter ($TRO$) and mid-toe ($MidToe$) markers (NOTE: this $\theta$ differs slightly from $\theta$ values calculated in leg stiffness above using Coleman et al. 2012).

$$\theta_{IC,MS} = atan2(TRO_Y - MidToe_Y, TRO_X - MidToe_X) - 90 * \frac{180}{\pi} \tag{10}$$

$$\theta = \theta_{MS} - \theta_{IC} \tag{11}$$

*10.1. Dimensionless stiffness calculations*

**Dimensionless leg compression**

$$\Delta L_{dim} = \frac{\Delta L}{L_0} \tag{12}$$

$L_0$ = leg length from greater trochanter to floor without shoes (m)

**Dimensionless resultant force**

$$F_{leg_{dim}} = \frac{F_{leg}}{BW} \tag{13}$$

$BW$ = body weight (kN)

**Dimensionless leg stiffness**

$$K_{LEG} = k_{leg} * \frac{L_0}{Mass} \tag{14}$$

**Dimensionless COM fluctuation**

$$\Delta y_{dim} = \frac{\Delta y}{L_0} \tag{15}$$

**Dimensionless max vertical force**

$$F_{V_{dim}} = \frac{F_V}{BW} \tag{16}$$

**Dimensionless vertical stiffness**

$$K_{VERT} = k_{vert} * \frac{L_0}{Mass} \tag{17}$$

### 11. Model results

Table A2: Slope estimates and model results for log(Mass) for Slow Froude and Fast Froude running. OLS = ordinary least squares regression; LMM = linear mixed model; BLMM = Bayesian linear mixed model. CI is 95% confidence interval/HDI.

| Variable | Speed | Model | Slope Estimate | 95% CI/HDI | $R^2$ |
|---|---|---|---|---|---|
| $F_{leg}$ | Slow Froude | OLS | 1.00 | 0.88, 1.11 | 0.82 |
| | | LMM | 1.00 | 0.88, 1.11 | 0.97 |
| | | BLMM | 0.99 | 0.88, 1.11 | 0.97 |
| | Fast Froude | OLS | 1.00 | 0.87, 1.12 | 0.81 |
| | | LMM | 1.00 | 0.88, 1.12 | 0.97 |
| | | BLMM | 1.00 | 0.88, 1.12 | 0.97 |
| $\Delta L$ | Slow Froude | OLS | 0.46 | 0.29, 0.63 | 0.32 |
| | | LMM | 0.46 | 0.29, 0.63 | 0.87 |
| | | BLMM | 0.46 | 0.29, 0.63 | 0.86 |
| | Fast Froude | OLS | 0.45 | 0.29, 0.61 | 0.34 |
| | | LMM | 0.45 | 0.29, 0.60 | 0.84 |
| | | BLMM | 0.45 | 0.29, 0.61 | 0.84 |
| $k_{leg}$ | Slow Froude | OLS | 0.53 | 0.32, 0.75 | 0.29 |
| | | LMM | 0.54 | 0.33, 0.75 | 0.88 |
| | | BLMM | 0.53 | 0.32, 0.75 | 0.88 |
| | Fast Froude | OLS | 0.55 | 0.35, 0.75 | 0.33 |
| | | LMM | 0.55 | 0.35, 0.74 | 0.85 |
| | | BLMM | 0.55 | 0.35, 0.75 | 0.85 |
| $F_V$ | Slow Froude | OLS | 1.00 | 0.88, 1.12 | 0.82 |
| | | LMM | 1.00 | 0.88, 1.11 | 0.97 |
| | | BLMM | 1.00 | 0.88, 1.12 | 0.97 |
| | Fast Froude | OLS | 1.00 | 0.87, 1.12 | 0.81 |
| | | LMM | 1.00 | 0.88, 1.12 | 0.97 |
| | | BLMM | 1.00 | 0.87, 1.11 | 0.97 |
| $\Delta y$ | Slow Froude | OLS | 0.39 | 0.19, 0.59 | 0.19 |
| | | LMM | 0.39 | 0.19, 0.59 | 0.77 |
| | | BLMM | 0.38 | 0.18, 0.59 | 0.77 |
| | Fast Froude | OLS | 0.37 | 0.14, 0.60 | 0.15 |
| | | LMM | 0.37 | 0.14, 0.59 | 0.80 |
| | | BLMM | 0.37 | 0.14, 0.59 | 0.80 |
| $k_{vert}$ | Slow Froude | OLS | 0.61 | 0.42, 0.80 | 0.40 |
| | | LMM | 0.61 | 0.43, 0.80 | 0.83 |
| | | BLMM | 0.61 | 0.42, 0.79 | 0.82 |
| | Fast Froude | OLS | 0.62 | 0.41, 0.84 | 0.37 |
| | | LMM | 0.63 | 0.42, 0.83 | 0.82 |
| | | BLMM | 0.63 | 0.42, 0.83 | 0.82 |
| $\theta$ | Slow Froude | OLS | 0.04 | -0.09, 0.17 | 0.01 |
| | | LMM | 0.04 | -0.09, 0.17 | 0.61 |
| | | BLMM | 0.04 | -0.09, 0.17 | 0.61 |
| | Fast Froude | OLS | 0.05 | -0.05, 0.16 | 0.02 |
| | | LMM | 0.05 | -0.05, 0.16 | 0.54 |
| | | BLMM | 0.06 | -0.05, 0.16 | 0.54 |

*Prior summary*

Shown below are default priors for linear mixed models before auto-scaling:

```
Priors for model '...'
------
Intercept (after predictors centered)
 ~ normal(location = 0, scale = 10)

Coefficients
 ~ normal(location = [0,0], scale = [2.5,2.5])

Auxiliary (sigma)
 ~ exponential(rate = 1)

Covariance
 ~ decov(reg. = 1, conc. = 1, shape = 1, scale = 1)
------
See help('prior_summary.stanreg') for more details
```

See Gabry and Goodrich (2019) for prior descriptions.

#### Full Bayesian model fit statistics

The following fit statistics are reported for the Bayesian models:

**Median** Median of the posterior distribution.

**HDI** 95% highest posterior density interval.

**ROPE, ROPE %** Region that is "practically equivalent" to zero, calculated from $[-0.1 * SD_y, 0.1 * SD_y]$, where $SD_y$ is the standard deviation of the response variable. ROPE % is the percentage of the 95% HDI that falls within the ROPE. 0% in the ROPE is considered a credibly non-zero parameter.

**Rhat** Measures the ratio of the average variance of the draws within each chain to the variance of the pooled draws across chains. Should not exceed 1.1.

**ESS** Effective sample size, or estimate of the effective number of independent draws from the posterior distribution of the parameter of interest. Because the draws within a chain are not independent if there is autocorrelation, the effective sample size will be smaller than the total number of iterations. Will also be reduced if samples are thinned.

**MCSE** Monte Carlo error, or the standard error of the *mean* of the posterior draws, is the uncertainty associated with the Monte Carlo approximation.

Descriptions from Kruschke (2014); Makowski et al. (2019); Muth et al. (2018).

Table A3: Bayesian model summary and fit statistics, including prior scales. Model parameters include: model intercept (Intercept) and slope (scaling exponent) for mass (log(Mass)).

| Model | Speed | Parameter | Median | 95% HDI | ROPE | ROPE % | Rhat | ESS | MCSE | Prior Scale |
|---|---|---|---|---|---|---|---|---|---|---|
| $F_{leg}$ | Slow | (Intercept) | -3.80 | -4.29, -3.32 | -0.02, 0.02 | 0.00 | 1.00 | 3378 | 0.00 | 1.96 |
| | | log(Mass) | 0.99 | 0.88, 1.12 | -0.02, 0.02 | 0.00 | 1.00 | 3380 | 0.00 | 2.81 |
| | Fast | (Intercept) | -3.72 | -4.22, -3.23 | -0.02, 0.02 | 0.00 | 1.00 | 2520 | 0.01 | 1.96 |
| | | log(Mass) | 1.00 | 0.88, 1.12 | -0.02, 0.02 | 0.00 | 1.00 | 2533 | 0.00 | 2.78 |
| $\Delta L$ | Slow | (Intercept) | -4.09 | -4.78, -3.37 | -0.02, 0.02 | 0.00 | 1.00 | 4197 | 0.01 | 1.53 |
| | | log(Mass) | 0.46 | 0.29, 0.63 | -0.02, 0.02 | 0.00 | 1.00 | 4181 | 0.00 | 2.20 |
| | Fast | (Intercept) | -4.00 | -4.67, -3.36 | -0.01, 0.01 | 0.00 | 1.00 | 4196 | 0.01 | 1.47 |
| | | log(Mass) | 0.45 | 0.29, 0.60 | -0.01, 0.01 | 0.00 | 1.00 | 4191 | 0.00 | 2.09 |
| $k_{leg}$ | Slow | (Intercept) | 0.30 | -0.60, 1.16 | -0.02, 0.02 | 2.86 | 1.00 | 2592 | 0.01 | 1.87 |
| | | log(Mass) | 0.53 | 0.32, 0.75 | -0.02, 0.02 | 0.00 | 1.00 | 2602 | 0.00 | 2.69 |
| | Fast | (Intercept) | 0.29 | -0.49, 1.14 | -0.02, 0.02 | 2.82 | 1.00 | 5396 | 0.01 | 1.81 |
| | | log(Mass) | 0.55 | 0.35, 0.74 | -0.02, 0.02 | 0.00 | 1.00 | 5399 | 0.00 | 2.57 |
| $F_V$ | Slow | (Intercept) | -3.83 | -4.29, -3.32 | -0.02, 0.02 | 0.00 | 1.00 | 3699 | 0.00 | 1.97 |
| | | log(Mass) | 1.00 | 0.88, 1.12 | -0.02, 0.02 | 0.00 | 1.00 | 3693 | 0.00 | 2.82 |
| | Fast | (Intercept) | -3.71 | -4.20, -3.21 | -0.02, 0.02 | 0.00 | 1.00 | 6168 | 0.00 | 1.96 |
| | | log(Mass) | 1.00 | 0.87, 1.11 | -0.02, 0.02 | 0.00 | 1.00 | 6151 | 0.00 | 2.78 |
| $\Delta y$ | Slow | (Intercept) | -4.29 | -5.16, -3.50 | -0.02, 0.02 | 0.00 | 1.00 | 2167 | 0.01 | 1.81 |
| | | log(Mass) | 0.38 | 0.20, 0.60 | -0.02, 0.02 | 0.00 | 1.00 | 2163 | 0.00 | 2.59 |
| | Fast | (Intercept) | -4.44 | -5.37, -3.52 | -0.02, 0.02 | 0.00 | 1.00 | 4069 | 0.01 | 1.89 |
| | | log(Mass) | 0.37 | 0.14, 0.59 | -0.02, 0.02 | 0.00 | 1.00 | 4037 | 0.00 | 2.67 |
| $k_{vert}$ | Slow | (Intercept) | 0.49 | -0.28, 1.25 | -0.02, 0.02 | 1.92 | 1.00 | 2445 | 0.01 | 1.85 |
| | | log(Mass) | 0.61 | 0.42, 0.79 | -0.02, 0.02 | 0.00 | 1.00 | 2443 | 0.00 | 2.65 |
| | Fast | (Intercept) | 0.73 | -0.13, 1.58 | -0.02, 0.02 | 0.92 | 1.00 | 4978 | 0.01 | 2.01 |
| | | log(Mass) | 0.63 | 0.42, 0.83 | -0.02, 0.02 | 0.00 | 1.00 | 4974 | 0.00 | 2.84 |
| $\theta$ | Slow | (Intercept) | 2.87 | 2.33, 3.42 | -0.01, 0.01 | 0.00 | 1.00 | 2344 | 0.01 | 1.16 |
| | | log(Mass) | 0.04 | -0.09, 0.17 | -0.01, 0.01 | 11.97 | 1.00 | 2341 | 0.00 | 1.67 |
| | Fast | (Intercept) | 2.97 | 2.56, 3.44 | -0.01, 0.01 | 0.00 | 1.00 | 4034 | 0.00 | 1.04 |
| | | log(Mass) | 0.06 | -0.06, 0.16 | -0.01, 0.01 | 9.54 | 1.00 | 4025 | 0.00 | 1.47 |

### Dimensionless model results

Table A4: Dimensionless variable slope estimates and model results for each model comparison

| Variable | Speed | Model | Slope Est | 95% CI/HDI | $R^2$ |
|---|---|---|---|---|---|
| $F_{leg_{dim}}$ | Slow Froude | OLS | 0.00 | -0.12, 0.11 | 0.00 |
| | | LMM | 0.00 | -0.12, 0.11 | 0.85 |
| | | BLMM | 0.00 | -0.12, 0.11 | 0.85 |
| | Fast Froude | OLS | 0.00 | -0.13, 0.12 | 0.00 |
| | | LMM | 0.00 | -0.12, 0.12 | 0.88 |
| | | BLMM | 0.00 | -0.12, 0.11 | 0.88 |
| $\Delta L_{dim}$ | Slow Froude | OLS | 0.02 | -0.14, 0.19 | 0.00 |
| | | LMM | 0.02 | -0.14, 0.18 | 0.81 |
| | | BLMM | 0.02 | -0.14, 0.18 | 0.80 |
| | Fast Froude | OLS | 0.01 | -0.14, 0.15 | 0.00 |
| | | LMM | 0.01 | -0.14, 0.15 | 0.76 |
| | | BLMM | 0.01 | -0.14, 0.16 | 0.76 |
| $K_{LEG}$ | Slow Froude | OLS | -0.03 | -0.25, 0.19 | 0.00 |
| | | LMM | -0.03 | -0.24, 0.19 | 0.85 |
| | | BLMM | -0.02 | -0.24, 0.19 | 0.84 |
| | Fast Froude | OLS | -0.01 | -0.21, 0.19 | 0.00 |
| | | LMM | -0.01 | -0.21, 0.18 | 0.79 |
| | | BLMM | -0.01 | -0.21, 0.19 | 0.79 |
| $F_{V_{dim}}$ | Slow Froude | OLS | 0.00 | -0.12, 0.12 | 0.00 |
| | | LMM | 0.00 | -0.12, 0.11 | 0.85 |
| | | BLMM | 0.00 | -0.12, 0.12 | 0.85 |
| | Fast Froude | OLS | 0.00 | -0.13, 0.12 | 0.00 |
| | | LMM | 0.00 | -0.12, 0.12 | 0.88 |
| | | BLMM | 0.00 | -0.12, 0.12 | 0.87 |
| $\Delta y_{dim}$ | Slow Froude | OLS | -0.04 | -0.25, 0.16 | 0.00 |
| | | LMM | -0.05 | -0.25, 0.16 | 0.75 |
| | | BLMM | -0.05 | -0.26, 0.15 | 0.74 |
| | Fast Froude | OLS | -0.07 | -0.30, 0.16 | 0.01 |
| | | LMM | -0.07 | -0.30, 0.16 | 0.78 |
| | | BLMM | -0.07 | -0.30, 0.16 | 0.78 |
| $K_{VERT}$ | Slow Froude | OLS | 0.05 | -0.16, 0.25 | 0.00 |
| | | LMM | 0.05 | -0.16, 0.25 | 0.78 |
| | | BLMM | 0.04 | -0.16, 0.25 | 0.77 |
| | Fast Froude | OLS | 0.06 | -0.16, 0.28 | 0.01 |
| | | LMM | 0.07 | -0.15, 0.28 | 0.76 |
| | | BLMM | 0.07 | -0.15, 0.29 | 0.76 |

*Dimensionless Bayesian model fit statistics*

Table A5: Bayesian model summary and fit statistics for dimensionless variables, including prior scales. Model parameters include: model intercept (Intercept) and coefficient (scaling exponent) for mass (log(Mass)).

| Model | Speed | Parameter | Median | 95% HDI | ROPE | ROPE % | Rhat | ESS | MCSE | Prior Scale |
|---|---|---|---|---|---|---|---|---|---|---|
| $F_{leg_{dim}}$ | Slow | (Intercept) | 0.81 | 0.34, 1.31 | -0.01, 0.01 | 0.00 | 1.00 | 3012 | 0.00 | 0.89 |
| | | log(Mass) | 0.00 | -0.12, 0.12 | -0.01, 0.01 | 13.12 | 1.00 | 3002 | 0.00 | 1.28 |
| | Fast | (Intercept) | 0.91 | 0.41, 1.39 | -0.01, 0.01 | 0.00 | 1.00 | 3135 | 0.00 | 0.90 |
| | | log(Mass) | 0.00 | -0.12, 0.11 | -0.01, 0.01 | 13.32 | 1.00 | 3102 | 0.00 | 1.27 |
| $\Delta L_{dim}$ | Slow | (Intercept) | -6.78 | -7.45, -6.11 | -0.01, 0.01 | 0.00 | 1.00 | 2354 | 0.01 | 1.26 |
| | | log(Mass) | 0.02 | -0.14, 0.18 | -0.01, 0.01 | 11.95 | 1.00 | 2334 | 0.00 | 1.82 |
| | Fast | (Intercept) | -6.66 | -7.27, -6.04 | -0.01, 0.01 | 0.00 | 1.00 | 3746 | 0.01 | 1.17 |
| | | log(Mass) | 0.01 | -0.14, 0.15 | -0.01, 0.01 | 13.29 | 1.00 | 3745 | 0.00 | 1.67 |
| $K_{LEG}$ | Slow | (Intercept) | 2.94 | 2.03, 3.80 | -0.02, 0.02 | 0.00 | 1.00 | 3474 | 0.01 | 1.67 |
| | | log(Mass) | -0.02 | -0.24, 0.19 | -0.02, 0.02 | 12.87 | 1.00 | 3460 | 0.00 | 2.40 |
| | Fast | (Intercept) | 2.95 | 2.11, 3.76 | -0.02, 0.02 | 0.00 | 1.00 | 3907 | 0.01 | 1.54 |
| | | log(Mass) | -0.01 | -0.20, 0.20 | -0.02, 0.02 | 12.99 | 1.00 | 3902 | 0.00 | 2.19 |
| $F_{V_{dim}}$ | Slow | (Intercept) | 0.80 | 0.32, 1.30 | -0.01, 0.01 | 0.00 | 1.00 | 4021 | 0.00 | 0.90 |
| | | log(Mass) | 0.00 | -0.12, 0.12 | -0.01, 0.01 | 12.65 | 1.00 | 4010 | 0.00 | 1.28 |
| | Fast | (Intercept) | 0.92 | 0.42, 1.42 | -0.01, 0.01 | 0.00 | 1.00 | 3733 | 0.00 | 0.89 |
| | | log(Mass) | 0.00 | -0.13, 0.11 | -0.01, 0.01 | 12.72 | 1.00 | 3755 | 0.00 | 1.27 |
| $\Delta y_{dim}$ | Slow | (Intercept) | -6.97 | -7.78, -6.09 | -0.02, 0.02 | 0.00 | 1.00 | 2209 | 0.01 | 1.69 |
| | | log(Mass) | -0.05 | -0.26, 0.15 | -0.02, 0.02 | 11.77 | 1.00 | 2209 | 0.00 | 2.41 |
| | Fast | (Intercept) | -7.08 | -8.03, -6.13 | -0.02, 0.02 | 0.00 | 1.00 | 3824 | 0.01 | 1.80 |
| | | log(Mass) | -0.07 | -0.30, 0.16 | -0.02, 0.02 | 10.59 | 1.00 | 3801 | 0.00 | 2.54 |
| $K_{VERT}$ | Slow | (Intercept) | 3.15 | 2.29, 3.99 | -0.02, 0.02 | 0.00 | 1.00 | 2528 | 0.01 | 1.62 |
| | | log(Mass) | 0.04 | -0.16, 0.25 | -0.02, 0.02 | 11.73 | 1.00 | 2541 | 0.00 | 2.32 |
| | Fast | (Intercept) | 3.38 | 2.51, 4.32 | -0.02, 0.02 | 0.00 | 1.00 | 4226 | 0.01 | 1.73 |
| | | log(Mass) | 0.07 | -0.16, 0.28 | -0.02, 0.02 | 10.47 | 1.00 | 4215 | 0.00 | 2.45 |

### Sex comparison Bayesian model fit statistics

Table A6: Bayesian model summary and fit statistics for sex interaction models. Model parameters include: model intercept (Intercept), coefficient (scaling exponent) for mass (log(Mass)), coefficient for male sex (SexMale), and interaction effect between mass and sex (log(Mass):SexMale).

| Model | Speed | Parameter | Median | 95% HDI | ROPE | ROPE % | Rhat | ESS | MCSE | Prior Scale |
|---|---|---|---|---|---|---|---|---|---|---|
| $F_{leg}$ | Slow | (Intercept) | -3.60 | -4.16, -2.97 | -0.02, 0.02 | 0.00 | 1.00 | 3046 | 0.01 | 1.96 |
| | | log(Mass) | 0.94 | 0.79, 1.09 | -0.02, 0.02 | 0.00 | 1.00 | 2984 | 0.00 | 2.81 |
| | | SexMale | 0.13 | -0.58, 0.75 | -0.02, 0.02 | 4.30 | 1.00 | 4197 | 0.01 | 0.49 |
| | | log(Mass):SexMale | -0.02 | -0.18, 0.14 | -0.02, 0.02 | 19.25 | 1.00 | 4179 | 0.00 | 0.23 |
| | Fast | (Intercept) | -3.34 | -3.91, -2.74 | -0.02, 0.02 | 0.00 | 1.00 | 3498 | 0.01 | 1.96 |
| | | log(Mass) | 0.90 | 0.75, 1.04 | -0.02, 0.02 | 0.00 | 1.00 | 3506 | 0.00 | 2.78 |
| | | SexMale | 0.02 | -0.61, 0.68 | -0.02, 0.02 | 4.99 | 1.00 | 5178 | 0.00 | 0.49 |
| | | log(Mass):SexMale | 0.01 | -0.14, 0.17 | -0.02, 0.02 | 20.17 | 1.00 | 5142 | 0.00 | 0.23 |
| $\Delta L$ | Slow | (Intercept) | -4.28 | -5.11, -3.46 | -0.02, 0.02 | 0.00 | 1.00 | 4716 | 0.01 | 1.53 |
| | | log(Mass) | 0.50 | 0.30, 0.71 | -0.02, 0.02 | 0.00 | 1.00 | 4725 | 0.00 | 2.20 |
| | | SexMale | 0.19 | -0.41, 0.81 | -0.02, 0.02 | 3.32 | 1.00 | 8238 | 0.00 | 0.38 |
| | | log(Mass):SexMale | -0.05 | -0.20, 0.10 | -0.02, 0.02 | 13.66 | 1.00 | 8172 | 0.00 | 0.18 |
| | Fast | (Intercept) | -3.88 | -4.62, -3.08 | -0.01, 0.01 | 0.00 | 1.00 | 4770 | 0.01 | 1.47 |
| | | log(Mass) | 0.41 | 0.22, 0.59 | -0.01, 0.01 | 0.00 | 1.00 | 4711 | 0.00 | 2.09 |
| | | SexMale | 0.27 | -0.31, 0.87 | -0.01, 0.01 | 2.60 | 1.00 | 8009 | 0.00 | 0.37 |
| | | log(Mass):SexMale | -0.05 | -0.20, 0.09 | -0.01, 0.01 | 13.29 | 1.00 | 7926 | 0.00 | 0.17 |
| $k_{leg}$ | Slow | (Intercept) | 0.71 | -0.32, 1.70 | -0.02, 0.02 | 1.17 | 1.00 | 4255 | 0.01 | 1.87 |
| | | log(Mass) | 0.43 | 0.18, 0.68 | -0.02, 0.02 | 0.00 | 1.00 | 4244 | 0.00 | 2.69 |
| | | SexMale | -0.13 | -0.88, 0.66 | -0.02, 0.02 | 3.74 | 1.00 | 7036 | 0.00 | 0.47 |
| | | log(Mass):SexMale | 0.05 | -0.14, 0.23 | -0.02, 0.02 | 14.91 | 1.00 | 6990 | 0.00 | 0.22 |
| | Fast | (Intercept) | 0.53 | -0.41, 1.50 | -0.02, 0.02 | 1.77 | 1.00 | 5342 | 0.01 | 1.81 |
| | | log(Mass) | 0.49 | 0.26, 0.72 | -0.02, 0.02 | 0.00 | 1.00 | 5294 | 0.00 | 2.57 |
| | | SexMale | -0.24 | -1.00, 0.45 | -0.02, 0.02 | 3.54 | 1.00 | 8454 | 0.00 | 0.45 |
| | | log(Mass):SexMale | 0.06 | -0.10, 0.25 | -0.02, 0.02 | 12.88 | 1.00 | 8418 | 0.00 | 0.21 |
| $F_V$ | Slow | (Intercept) | -3.60 | -4.18, -2.98 | -0.02, 0.02 | 0.00 | 1.00 | 3267 | 0.01 | 1.97 |
| | | log(Mass) | 0.94 | 0.79, 1.09 | -0.02, 0.02 | 0.00 | 1.00 | 3272 | 0.00 | 2.82 |
| | | SexMale | 0.13 | -0.49, 0.84 | -0.02, 0.02 | 4.79 | 1.00 | 4072 | 0.01 | 0.49 |
| | | log(Mass):SexMale | -0.02 | -0.18, 0.14 | -0.02, 0.02 | 20.00 | 1.00 | 4040 | 0.00 | 0.23 |
| | Fast | (Intercept) | -3.34 | -3.93, -2.76 | -0.02, 0.02 | 0.00 | 1.00 | 5449 | 0.00 | 1.96 |
| | | log(Mass) | 0.90 | 0.75, 1.04 | -0.02, 0.02 | 0.00 | 1.00 | 5440 | 0.00 | 2.78 |
| | | SexMale | 0.04 | -0.62, 0.69 | -0.02, 0.02 | 4.97 | 1.00 | 6549 | 0.00 | 0.49 |
| | | log(Mass):SexMale | 0.01 | -0.15, 0.17 | -0.02, 0.02 | 20.23 | 1.00 | 6511 | 0.00 | 0.23 |
| $\Delta y$ | Slow | (Intercept) | -4.88 | -5.84, -3.93 | -0.02, 0.02 | 0.00 | 1.00 | 3672 | 0.01 | 1.81 |
| | | log(Mass) | 0.54 | 0.30, 0.77 | -0.02, 0.02 | 0.00 | 1.00 | 3653 | 0.00 | 2.59 |
| | | SexMale | 0.12 | -0.59, 0.84 | -0.02, 0.02 | 4.06 | 1.00 | 7093 | 0.00 | 0.45 |
| | | log(Mass):SexMale | -0.05 | -0.22, 0.12 | -0.02, 0.02 | 14.17 | 1.00 | 7050 | 0.00 | 0.22 |
| | Fast | (Intercept) | -4.85 | -6.03, -3.79 | -0.02, 0.02 | 0.00 | 1.00 | 3460 | 0.01 | 1.89 |
| | | log(Mass) | 0.47 | 0.21, 0.76 | -0.02, 0.02 | 0.00 | 1.00 | 3458 | 0.00 | 2.67 |
| | | SexMale | 0.21 | -0.55, 0.96 | -0.02, 0.02 | 3.46 | 1.00 | 6873 | 0.00 | 0.47 |
| | | log(Mass):SexMale | -0.06 | -0.25, 0.12 | -0.02, 0.02 | 12.88 | 1.00 | 6796 | 0.00 | 0.22 |
| $k_{vert}$ | Slow | (Intercept) | 1.31 | 0.47, 2.11 | -0.02, 0.02 | 0.00 | 1.00 | 4636 | 0.01 | 1.85 |
| | | log(Mass) | 0.39 | 0.20, 0.60 | -0.02, 0.02 | 0.00 | 1.00 | 4637 | 0.00 | 2.65 |
| | | SexMale | -0.09 | -0.78, 0.62 | -0.02, 0.02 | 4.19 | 1.00 | 7509 | 0.00 | 0.46 |
| | | log(Mass):SexMale | 0.06 | -0.11, 0.22 | -0.02, 0.02 | 13.97 | 1.00 | 7463 | 0.00 | 0.22 |
| | Fast | (Intercept) | 1.56 | 0.63, 2.52 | -0.02, 0.02 | 0.00 | 1.00 | 8779 | 0.01 | 2.01 |
| | | log(Mass) | 0.41 | 0.18, 0.64 | -0.02, 0.02 | 0.00 | 1.00 | 8755 | 0.00 | 2.84 |
| | | SexMale | -0.27 | -1.06, 0.51 | -0.02, 0.02 | 3.28 | 1.00 | 10505 | 0.00 | 0.50 |
| | | log(Mass):SexMale | 0.10 | -0.09, 0.28 | -0.02, 0.02 | 10.83 | 1.00 | 10448 | 0.00 | 0.24 |

Table A6: Bayesian model summary and fit statistics for sex interaction models *(continued)*

| Model | Speed | Parameter | Median | 95% HDI | ROPE | ROPE % | Rhat | ESS | MCSE | Prior Scale |
|---|---|---|---|---|---|---|---|---|---|---|
| $\theta$ | Slow | (Intercept) | 2.82 | 2.21, 3.50 | -0.01, 0.01 | 0.00 | 1.00 | 3598 | 0.01 | 1.16 |
| | | log(Mass) | 0.05 | -0.11, 0.21 | -0.01, 0.01 | 10.01 | 1.00 | 3579 | 0.00 | 1.67 |
| | | SexMale | 0.01 | -0.46, 0.47 | -0.01, 0.01 | 4.15 | 1.00 | 7765 | 0.00 | 0.29 |
| | | log(Mass):SexMale | 0.00 | -0.12, 0.10 | -0.01, 0.01 | 17.16 | 1.00 | 7620 | 0.00 | 0.14 |
| | Fast | (Intercept) | 3.01 | 2.50, 3.55 | -0.01, 0.01 | 0.00 | 1.00 | 5282 | 0.00 | 1.04 |
| | | log(Mass) | 0.04 | -0.08, 0.18 | -0.01, 0.01 | 10.71 | 1.00 | 5287 | 0.00 | 1.47 |
| | | SexMale | 0.05 | -0.36, 0.46 | -0.01, 0.01 | 3.90 | 1.00 | 8810 | 0.00 | 0.26 |
| | | log(Mass):SexMale | -0.01 | -0.11, 0.09 | -0.01, 0.01 | 16.83 | 1.00 | 8562 | 0.00 | 0.12 |
